# Phosphate-responsive ornithine lipid biosynthesis remodels outer membrane composition and contributes to polymyxin tolerance

**DOI:** 10.64898/2026.11.03.764129

**Authors:** Roberto Jhonatan Olea-Ozuna, Melanie J. Campbell, Prithvijeet Singh Kharnal, Fabiha Zaheen Khan, Berenice Furlan, Samantha Y. Quintanilla, Sinjini Nandy, Kelli L. Palmer, Jennifer S. Brodbelt, Joseph M. Boll

## Abstract

Bacteria frequently encounter environments in which essential nutrients are limiting. Phosphate limitation constrains phospholipid synthesis and challenges maintenance of the cell envelope. While many Gram-negative bacteria respond by producing phosphorous-free membrane lipids, the mechanisms and consequences of this remodeling remain poorly understood in Enterobacterales. Here, we characterize the genetic and physiological basis of ornithine lipid (OL) biosynthesis in the opportunistic pathogens *Enterobacter cloacae* and *Klebsiella pneumoniae*. We show that OL biosynthesis depends on OlsF, a bifunctional acyltransferase that catalyzes both acylation steps required for OL synthesis. Phosphate limitation promotes OlsF-dependent OL accumulation through the PhoBR two-component system. Membrane fractionation revealed that OlsF-derived OLs are present in both membrane compartments but are enriched in the outer membrane. OlsF-dependent OL production enhances polymyxin tolerance during phosphate limitation, linking nutrient availability to changes in antimicrobial susceptibility. Together, these findings identify OlsF-dependent OL synthesis as a phosphate-responsive membrane remodeling pathway that reshapes outer membrane composition and influences the physiological properties of the Gram-negative cell envelope.

**Importance:** Bacteria remodel their membrane lipid composition when phosphate becomes limiting, but how this process influences the physiology of the Gram-negative cell envelope remains poorly understood. Here, we identify OlsF as the bifunctional enzyme responsible for phosphate-responsive ornithine lipid biosynthesis in the opportunistic pathogens *Enterobacter cloacae* and *Klebsiella pneumoniae*. We show that OlsF-derived ornithine lipids become enriched in the outer membrane and contribute to polymyxin tolerance during phosphate limitation. These findings reveal a phosphate-responsive membrane remodeling pathway that links nutrient availability to outer membrane lipid composition and antimicrobial susceptibility, highlighting how environmental conditions can reshape the functional properties of the Gram-negative cell envelope.

## Introduction

Phospholipids form the structural foundation of the inner and outer membranes of diderm bacteria and are central to membrane organization and function (1, 2). Because their synthesis requires inorganic phosphate (P_i_), changes in phosphate availability can influence membrane biogenesis and composition (2). In many environmental and host-associated niches, P_i_ availability can be limiting, requiring bacteria to maintain envelope integrity while conserving this essential nutrient (3, 4). One adaptation to phosphate limitation is the remodeling of membrane lipid composition through the production of phosphorus-free lipids, including glycolipids and aminolipids (5, 6). These adaptive responses can reshape envelope composition and influence how bacteria interact with environmental stresses, antimicrobial compounds, and host-associated niches (7–9).

Ornithine lipids (OLs) are widely distributed aminolipids whose production varies among bacterial species and environmental conditions (2, 10). OLs are produced constitutively in some bacteria, whereas in others their abundance changes in response to environmental cues, including phosphate availability (11, 12). Their biosynthesis involves sequential *N*-acylation of ornithine followed by *O*-acylation of the resulting mono-acylated intermediate to generate a di-acylated membrane lipid (13, 14). In many Proteobacteria, including *Pseudomonas aeruginosa* and *Acinetobacter baumannii*, these reactions are catalyzed by the two-enzyme OlsB/OlsA pathway (7, 9, 15), as illustrated in **Figure S1**. Alternative biosynthetic architectures have also been identified, including the bifunctional acyltransferase OlsF, first characterized in *Serratia proteamaculans* (16) and subsequently identified in other bacterial species, including *Vibrio cholerae* (17, 18), and the distinct OlsH/OlsI pathway identified in Planctomycetes (19). The identification of *olsF* homologs in several Enterobacterales lineages suggests that this biosynthetic pathway may extend beyond the organisms in which it was initially characterized (20). However, whether OlsF drives phosphate-dependent membrane remodeling in Enterobacterales, and how this remodeling influences cell envelope biology, remains poorly understood.

Beyond serving as phosphorus-free alternatives to phospholipids, OLs have the potential to alter the physicochemical properties of the bacterial envelope (7, 10). This is particularly relevant to the outer membrane, which serves as the primary permeability barrier of Gram-negative bacteria and provides protection against environmental stresses and membrane-targeting antimicrobials, including polymyxins (1, 21). Changes in outer membrane lipid composition can influence barrier function and antimicrobial susceptibility (22, 23). Studies in *P. aeruginosa*, *A. baumannii*, and *V. cholerae* have linked phosphorus-free lipid production to changes in antimicrobial susceptibility or tolerance (7–9, 18). However, whether phosphate-responsive OlsF-dependent OL production influences membrane organization and polymyxin tolerance in clinically important Enterobacterales remains poorly understood. In particular, the distribution of OlsF-derived OLs between the inner and outer membranes has not been established in these organisms. These gaps limit our understanding of how nutrient availability influences membrane composition and antimicrobial responses.

Here, we investigated the regulation, membrane distribution, and physiological role of OlsF-dependent OL biosynthesis in *Enterobacter cloacae* and *Klebsiella pneumoniae*. We show that OlsF-dependent OL synthesis is associated with the PhoBR-mediated phosphate limitation response, that OLs become enriched in the outer membrane, and that their production contributes to polymyxin tolerance. Together, these findings establish OlsF-dependent OL synthesis as a phosphate-responsive membrane remodeling pathway that links nutrient availability to outer membrane composition and antimicrobial tolerance.

## Results

### Conserved OlsF homologs direct ornithine lipid biosynthesis in Enterobacterales

Analysis of previously published datasets (20) revealed that OlsF homologs are common in several Enterobacterales lineages and were identified in all examined *Enterobacter* (21/21) and *Klebsiella* (11/11) genomes, and most *Serratia* genomes (16/17), but were not identified in representative *Escherichia*, *Shigella*, *Salmonella*, or *Yersinia* genomes under the applied search criteria (**Figure 1A**; **Table S1**). Homologs of OlsB and OlsH, proteins associated with alternative OL biosynthetic pathways, were also not identified among the examined genomes under the same search criteria. These observations suggest that OlsF-dependent OL synthesis is a conserved feature of specific Enterobacterales lineages, including clinically important *Enterobacter* and *Klebsiella* species. The widespread conservation of OlsF within these genera suggests that phosphate-responsive OL production may constitute a conserved membrane adaptation strategy among clinically important Enterobacterales.

**Figure 1.**
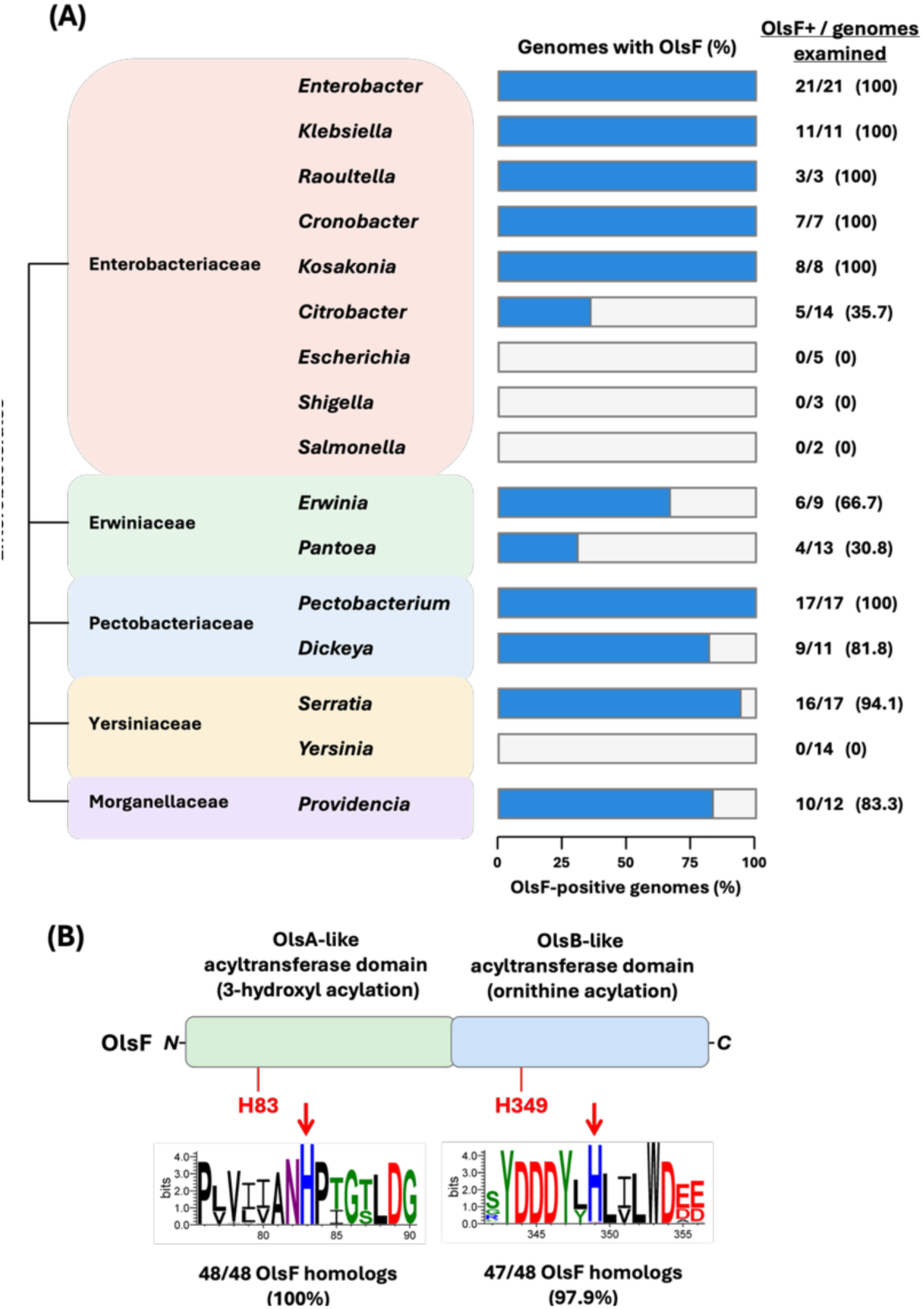
Distribution, domain architecture, and sequence conservation of OlsF in Enterobacterales. **(A)** Distribution of OlsF homologs among reference genomes from selected Enterobacterales genera based on a focused analysis of previously reported genomic datasets. Bars indicate the percentage of examined genomes containing an OlsF homolog, with the number of OlsF-positive genomes relative to the total number examined shown for each genus. Genera are grouped by taxonomic family. **(B)** Predicted domain architecture of OlsF and conservation of the regions surrounding H83 and H349. Sequence logos were generated from an alignment of 48 unique OlsF homologs from *Enterobacter*, *Klebsiella*, and *Serratia*. Residue numbering corresponds to OlsF from *Serratia proteamaculans* (WP_099062946.1). The number and percentage of homologs retaining histidine at the positions corresponding to H83 and H349 are indicated below the respective sequence logos.

OlsF is a single polypeptide that catalyzes both acyltransferase reactions performed by OlsA and OlsB in the canonical two-enzyme pathway. Consistent with the previously characterized OlsF from *S. proteamaculans* (16), the OlsF homologs from *E. cloacae* and *K. pneumoniae* contain predicted *N*-terminal OlsA-like and *C*-terminal OlsB-like regions (**Figure 1B**). To identify candidate catalytic residues conserved across Enterobacterales OlsF proteins, sequence comparison of 48 OlsF homologs from *Enterobacter*, *Klebsiella*, and *Serratia* revealed strong conservation of two histidine residues located within these respective regions. The position corresponding to H83 of *S. proteamaculans* OlsF was invariant across all 48 sequences, whereas H349 was conserved in 47 of 48 homologs (97.9%) (**Figure 1B**).

To experimentally test whether OlsF alone is sufficient for OL biosynthesis, we heterologously expressed each *olsF* allele in *Escherichia coli* W3110, which lacks known OL biosynthetic genes and does not naturally produce detectable OLs. Expression of either allele resulted in the appearance of a distinct ninhydrin-reactive aminolipid detectable by thin-layer chromatography (**Figure 2A**), in addition to production of multiple classes of lipids, including phosphatidylethanolamines (PE), modified PEs, and cardiolipins (CL) as detected in the MS1 spectra (**Figure 2B**). Specific lipid fractions were recovered from thin-layer chromatography plates 2 (**Figure S2**) and analyzed by shotgun mass spectrometry. The lipid profiles of the two OlsF samples are shown in **Figure 2B**, and tandem mass spectrometry (MS/MS) was used to confirm the structural assignments as summarized in **Table S2**. The MS/MS analysis confirmed the presence of OLs based on the characteristic loss of the ornithine headgroup, yielding the consistent fragment ion of *m/z* 131.08 (**Figure S3A**). In addition, a C_2_H_3_O_2_-modified OL of *m/z* 681 was also identified for both OlsF samples based on its characteristic fragmentation pattern (**Figure S3B**). For the W3110 + OlsF*_Ecl_* sample, isolation and activation of the ion peak of *m/z* 681.55 (**Figure 2B**) produced a chimeric MS/MS spectrum containing fragment ions from both the putative C_2_H_3_O_2_-modified OL 16:0/16:0 and CL 16:0_17:1/16:0_16:0 (**Figure S3B**).

**Figure 2.**
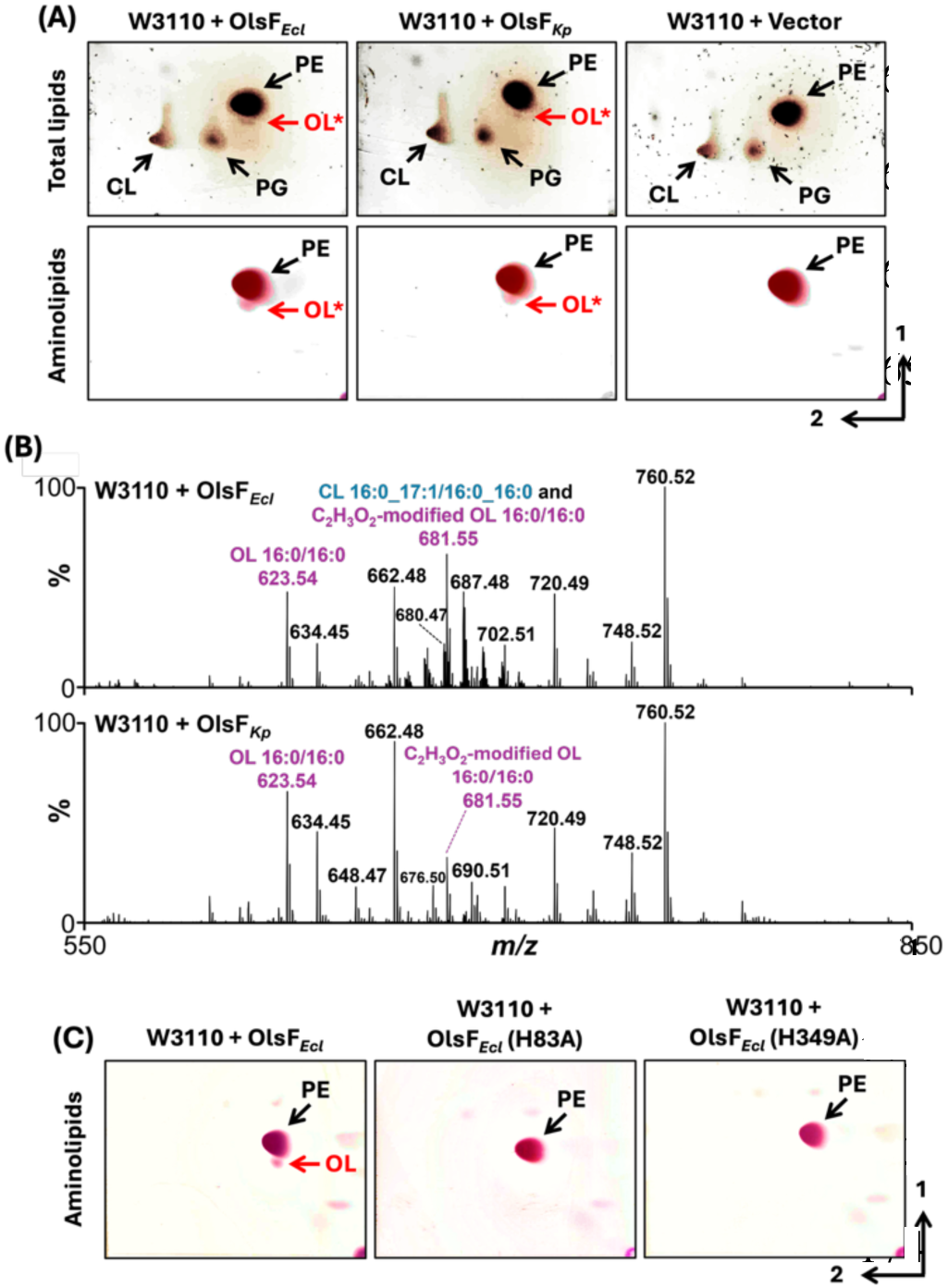
Identification of ornithine lipids produced by heterologous expression of OlsF in *E. coli*. **(A)** Representative two-dimensional thin-layer chromatography (TLC) analysis of total lipids (upper panels) and aminolipids (lower panels) extracted from *E. coli* W3110 expressing OlsF from *E. cloacae* (OlsF*_Ecl_*) or *K. pneumoniae* (OlsF*_Kp_*) or carrying the empty vector. Total lipids and aminolipids were visualized using sulfuric acid and ninhydrin staining, respectively. **(B)** Representative mass spectra of lipid fractions recovered from *E. coli* W3110 expressing OlsF*_Ecl_* or OlsF*_Kp_*. Ion peaks corresponding to putative ornithine lipid (OL) species are highlighted in purple, including ions of *m/z* 623.54 and 681.55. Proposed lipid assignments are indicated above the corresponding peaks. Spectra were obtained from lipid fractions recovered from the OL-containing regions of TLC plates shown in **Figure S2**. **(C)** Representative two-dimensional TLC analysis of aminolipids extracted from *E. coli* W3110 expressing wild-type OlsF*_Ecl_* or the H83A and H349A variants. OL, ornithine lipid; PE, phosphatidylethanolamine; PG, phosphatidylglycerol; CL, cardiolipin. Red arrows indicate putative OL species detected following heterologous OlsF expression.

We next examined whether the conserved histidine residues identified by sequence analysis were required for detectable OL production. H83A and H349A substitutions were introduced individually into the *E. cloacae olsF* allele and expressed in *E. coli* W3110. In contrast to wild-type OlsF, neither variant produced detectable OL under the conditions tested (**Figure 2C**). These findings demonstrate that two highly conserved histidine residues (H83 and H349) are essential for OlsF-dependent OL biosynthesis, consistent with conserved catalytic roles across Enterobacterales OlsF homologs.

### OlsF is required for phosphate-responsive ornithine lipid accumulation

Our previous work showed that *E. cloacae* and *K. pneumoniae* accumulate OLs under phosphate-limiting conditions (9). We therefore asked whether OlsF is required for this response under native conditions. Cells were grown under phosphate-replete (1 mM P_i_) and phosphate-limiting (50 µM P_i_) conditions, and growth was monitored under both conditions (**Figure S4A**). Consistent with our previous findings, OLs were not detected under phosphate-replete conditions but accumulated in both species during phosphate limitation (**Figure 3**). Quantification of the thin-layer chromatography profiles revealed a marked shift in membrane lipid composition under phosphate limitation, characterized by the accumulation of OLs alongside the major phospholipids phosphatidylethanolamine (PE), phosphatidylglycerol (PG), and cardiolipin (CL) (**Table S3**). These findings indicate that phosphate limitation promotes substantial membrane lipid remodeling in both species. We next examined whether phosphate limitation was accompanied by changes in lipopolysaccharide profiles. SDS-PAGE analysis revealed no major qualitative differences in lipopolysaccharide migration patterns between phosphate-replete and phosphate-limiting conditions in either species (**Figure S4B**).

**Figure 3.**
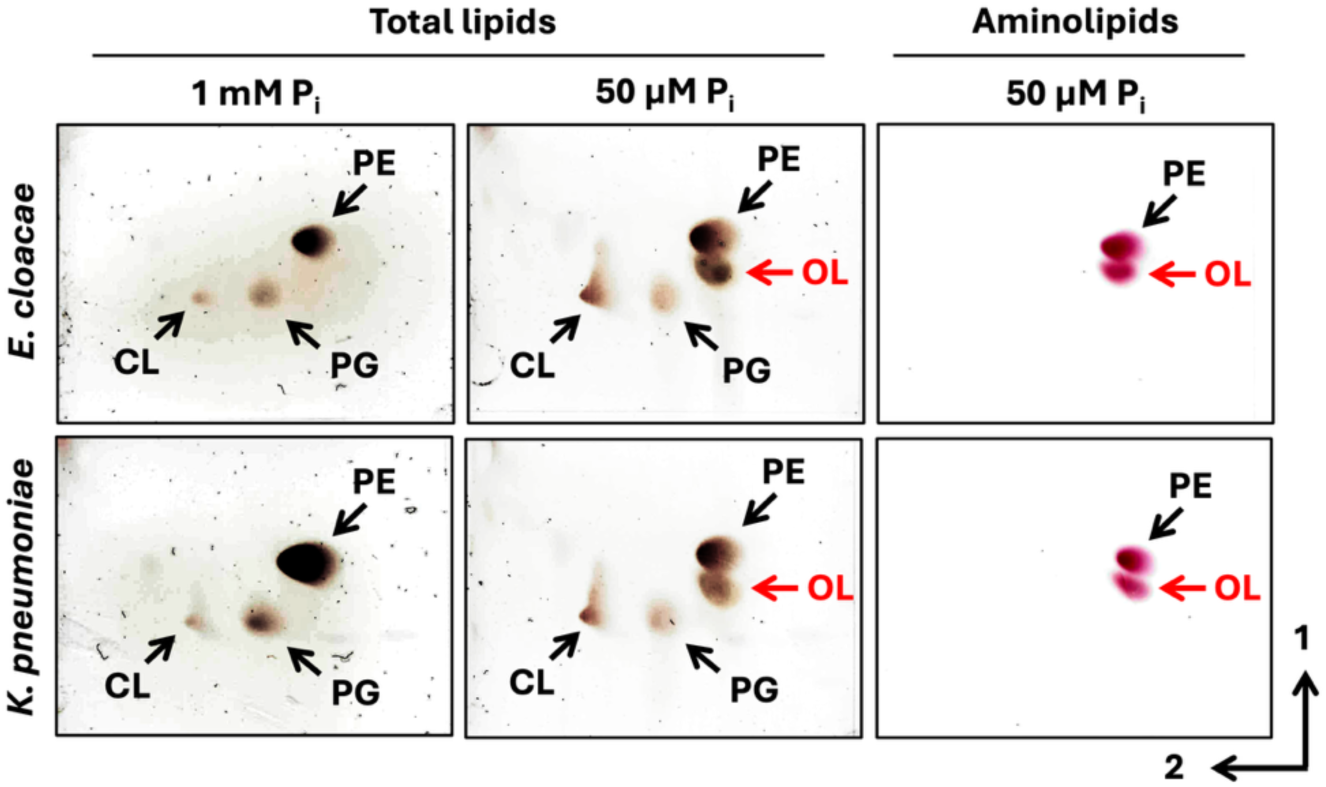
Phosphate limitation promotes membrane lipid remodeling and ornithine lipid accumulation in *E. cloacae* and *K. pneumoniae*. Two-dimensional thin-layer chromatography (TLC) analysis of total lipids extracted from *E. cloacae* and *K. pneumoniae* grown under phosphate-replete (1 mM Pi) or phosphate-limiting (50 µM Pi) conditions. Total lipids were visualized by sulfuric acid staining, whereas aminolipids were detected by ninhydrin staining. OLs accumulated under phosphate-limiting conditions in both species but were not detected under phosphate-replete conditions. Arrows indicate phosphatidylethanolamine (PE), phosphatidylglycerol (PG), cardiolipin (CL), and ornithine lipids (OLs). Numbers indicate the first and second dimensions of TLC separation.

To directly test the requirement for OlsF, we generated an in-frame Δ*olsF* mutant in *E. cloacae*. The Δ*olsF* strain failed to accumulate detectable OLs under phosphate-limiting conditions, whereas complementation restored OL production (**Figure 4A**). Thus, OlsF is essential for phosphate-responsive OL accumulation and the membrane lipid remodeling observed during phosphate limitation.

**Figure 4.**
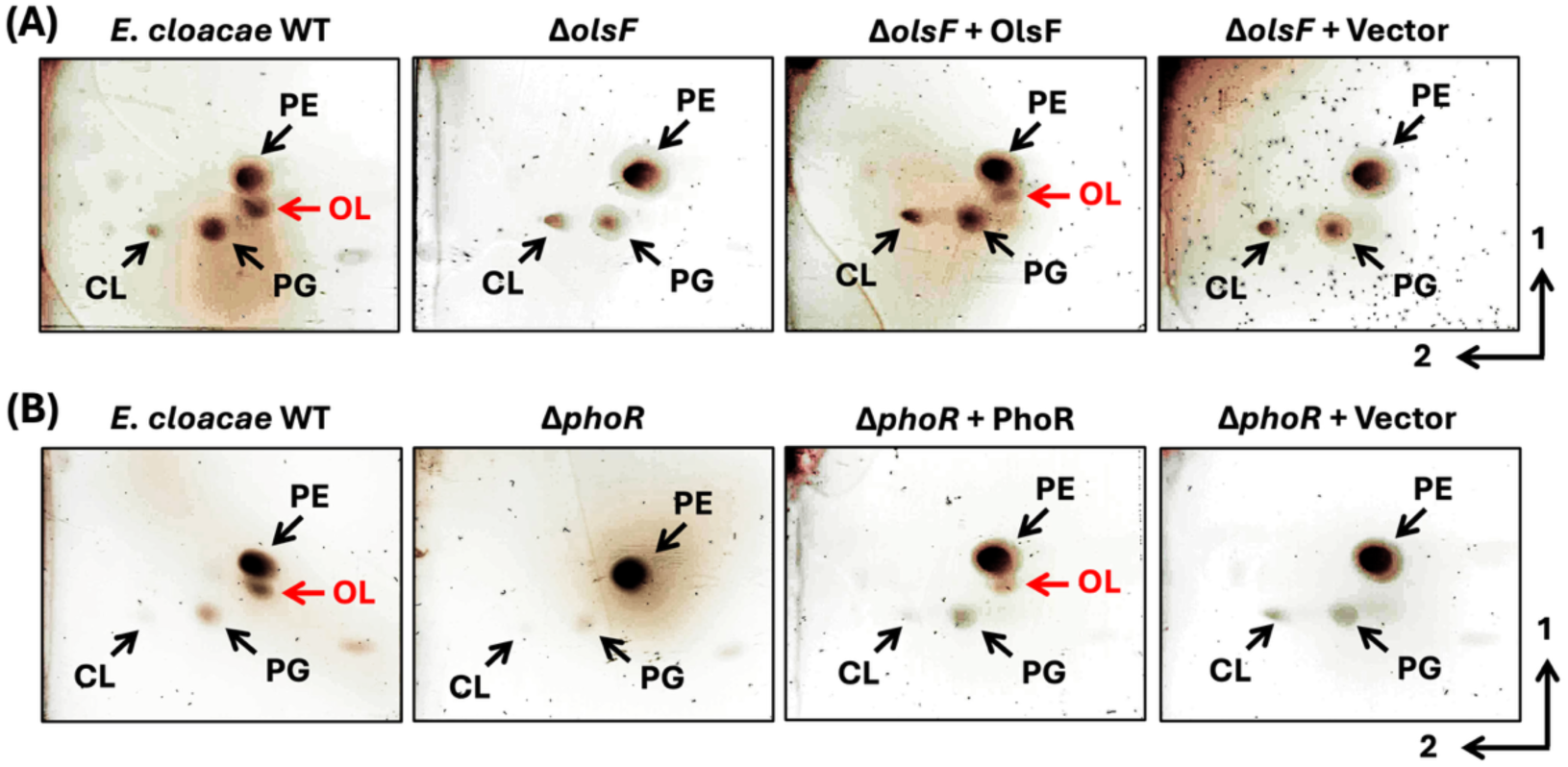
OlsF and PhoR are required for ornithine lipid accumulation during phosphate limitation in *E. cloacae*. Two-dimensional thin-layer chromatography (TLC) analysis of total lipids extracted from *E. cloacae* strains grown under phosphate-limiting conditions (50 µM P_i_). **(A)** Lipid profiles of wild-type (WT), Δ*olsF*, Δ*olsF* complemented with *olsF*, and Δ*olsF* carrying the empty vector. Ornithine lipids (OLs) were detected in the WT and complemented strains but were absent from the Δ*olsF* mutant and vector control. **(B)** Lipid profiles of WT, Δ*phoR*, Δ*phoR* complemented with *phoR*, and Δ*phoR* carrying the empty vector. OLs were detected in the WT and complemented strains but were absent from the Δ*phoR* mutant and vector control. Lipids were visualized by sulfuric acid staining. Arrows indicate phosphatidylethanolamine (PE), phosphatidylglycerol (PG), cardiolipin (CL), and ornithine lipids (OLs). Numbers indicate the first and second dimensions of TLC separation.

Phosphate limitation also induced moderate cell elongation in both *E. cloacae* and *K. pneumoniae* relative to phosphate-replete conditions (**Figure S4C and S4D**), consistent with what has been observed in other Gram-negative bacteria (6, 9). However, loss of OL synthesis did not further alter cell morphology or growth kinetics under phosphate-limiting conditions, as the Δ*olsF* mutant exhibited phenotypes comparable to those of the wild-type strain (**Figure S4E–G**). These observations indicate that OlsF-dependent OL production is not required for growth under phosphate limitation but instead may contribute to more specialized aspects of envelope adaptation.

### PhoR is required for phosphate-responsive ornithine lipid biosynthesis

Because the PhoBR two-component system coordinates bacterial responses to phosphate limitation (3, 24), we investigated whether it contributes to OL accumulation in *E. cloacae*. To test this, we examined OL synthesis in an in-frame Δ*phoR* mutant of *E. cloacae*. In contrast to the wild-type strain, the Δ*phoR* mutant failed to accumulate OLs under phosphate-limiting conditions (**Figure 4B**). Complementation with *phoR* in *trans* restored OL production (**Figure 4B**), demonstrating that PhoR is required for OlsF-dependent OL synthesis during phosphate limitation. These findings place OlsF-dependent OL production within the canonical PhoBR phosphate starvation response and identify phosphate sensing as a key regulator of membrane lipid remodeling in *E. cloacae*.

Analysis of the upstream regions of *olsF* in *E. cloacae* and *K. pneumoniae* revealed sequences resembling the consensus PhoB-binding motif (**Figure S5A**). The presence of these motifs, together with the loss of OL production in the Δ*phoR* mutant, supports a model in which OlsF-dependent OL biosynthesis is incorporated into the PhoBR regulatory network that coordinates adaptation to phosphate limitation.

Consistent with the established PhoBR-dependent regulation of the OlsB/OlsA pathway in *P. aeruginosa*, OL accumulation was also abolished in PA14 *phoB* and *phoR* transposon mutants under phosphate-limiting conditions (**Figure S5B**).

Together, these observations suggest that PhoBR-mediated control of OL biosynthesis may represent a conserved feature of phosphate-responsive membrane remodeling across diverse Gram-negative bacteria. Collectively, these findings establish PhoBR as a key regulator of phosphate-responsive OL production and link environmental phosphate sensing to membrane lipid remodeling in Enterobacterales.

### OlsF-dependent membrane remodeling enhances polymyxin tolerance during phosphate limitation

Because membrane lipid composition can influence outer membrane function and susceptibility to membrane-targeting antibiotics, we investigated whether OlsF-dependent OL production contributes to polymyxin tolerance. We first assessed the susceptibility of *E. cloacae* and *K. pneumoniae* to polymyxin B and colistin under phosphate-replete and phosphate-limiting conditions. Both species exhibited smaller inhibition zones under phosphate limitation than under phosphate-replete conditions (**Table S4**). Consistent with these observations, survival assays revealed increased survival following polymyxin B exposure in both species under phosphate-limiting conditions (**Figure 5A**). These findings indicate that phosphate limitation promotes a physiological state associated with enhanced polymyxin tolerance.

**Figure 5.**
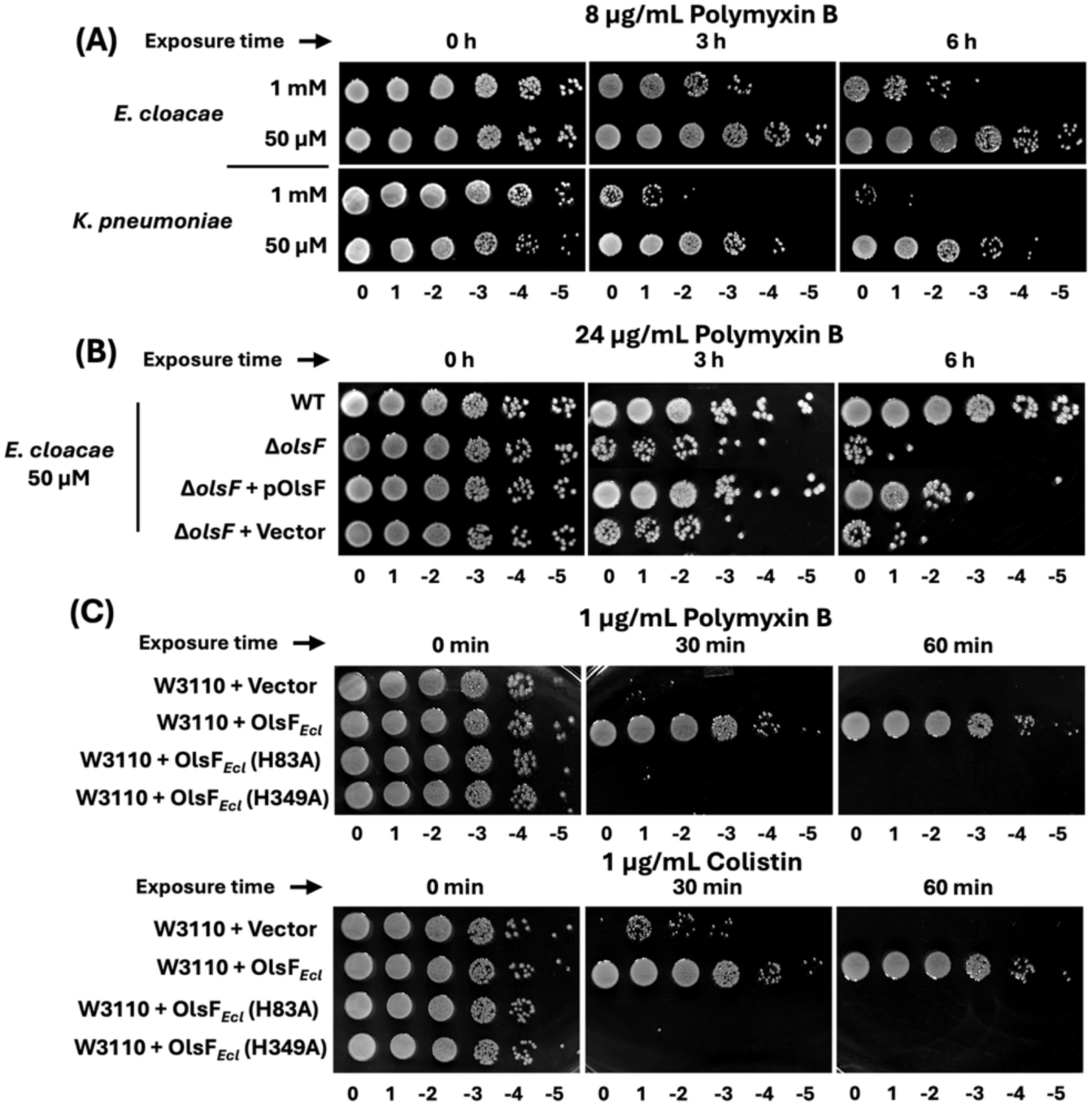
OlsF-dependent ornithine lipid production contributes to polymyxin tolerance. **(A)** Survival of *E. cloacae* and *K. pneumoniae* grown under phosphate-replete (1 mM P_i_) or phosphate-limiting (50 µM P_i_) conditions following exposure to 8 µg mL⁻¹ polymyxin B for 0, 3, and 6 h. Both species exhibited increased survival under phosphate-limiting conditions relative to phosphate-replete conditions. **(B)** Survival of *E. cloacae* wild-type (WT), Δ*olsF*, Δ*olsF* complemented with *olsF*, and Δ*olsF* carrying the empty vector under phosphate-limiting conditions (50 µM Pi) following exposure to 24 µg mL⁻¹ polymyxin B for 0, 3, and 6 h. Loss of *olsF* reduced polymyxin tolerance, whereas complementation restored the phenotype. **(C)** Survival of *E. coli* W3110 carrying the empty vector, expressing wild-type *E. cloacae* OlsF, or expressing the OlsF H83A or H349A variants following exposure to 1 µg mL⁻¹ polymyxin B or 1 µg mL⁻¹ colistin for 0, 30, and 60 min. Expression of wild-type OlsF increased survival following polymyxin exposure, whereas the catalytically inactive H83A and H349A variants did not. At each time point, cultures were serially diluted tenfold and spotted onto LB agar. Representative survival assays from three independent biological replicates are shown.

To determine whether OlsF-dependent OL production contributes to this phenotype, we examined the *E. cloacae* Δ*olsF* mutant and complemented strains under phosphate-limiting conditions using a higher concentration of polymyxin B to assess their response to increased antibiotic stress. Under these conditions, the Δ*olsF* mutant exhibited reduced survival compared with the wild-type strain, whereas complementation partially restored survival (**Figure 5B**). These results indicate that OlsF contributes to polymyxin tolerance during phosphate limitation.

We further investigated whether OlsF expression could enhance polymyxin tolerance in a heterologous host. Expression of wild-type *E. cloacae* OlsF in *E. coli* W3110 resulted in detectable OL production and increased survival following exposure to both polymyxin B and colistin compared with the empty-vector control (**Figure 5C**). In contrast, the OlsF H83A and H349A variants, which failed to produce detectable OLs, did not confer the survival advantage observed with wild-type OlsF (**Figure 5C**). Together, these findings establish a functional link between OlsF-dependent OL production and polymyxin tolerance, suggesting that phosphate-responsive membrane lipid remodeling contributes to adaptation against membrane-targeting antibiotics.

### Outer membrane enrichment of ornithine lipids provides a spatial basis for phosphate responsive membrane remodeling

Given the role of OlsF-dependent OL production to polymyxin tolerance and the central role of the outer membrane in interactions with membrane-targeting antibiotics, we next examined the membrane distribution of OlsF-derived OLs. Inner and outer membrane fractions were isolated from *E. cloacae* and *K. pneumoniae* grown under phosphate-limiting conditions, as well as from *E. coli* W3110 expressing *E. cloacae olsF* grown in LB. Separation of the membrane fractions was supported by enrichment of NADH oxidase activity in the inner membrane fractions and lipopolysaccharide in the outer membrane fractions (**Figure S6A–C**).

Lipid extraction followed by thin-layer chromatography revealed that OLs were present in both inner and outer membrane fractions of *E. cloacae* and *K. pneumoniae* (**Figure 6A**). Quantification of lipid composition showed that OLs accounted for a greater proportion of the total lipid signal in the outer membrane than in the inner membrane of both species (**Figure 6B and C**). Specifically, OLs represented approximately 33% and 22% of the lipid signal in the outer and inner membranes of *E. cloacae*, respectively, compared with approximately 37% and 26% in *K. pneumoniae*. The preferential accumulation of OLs in the outer membrane is notable given the role of this compartment in permeability barrier function and susceptibility to membrane-targeting antimicrobials. The relative proportions of the other major phospholipid classes also differed between membrane fractions (**Figure S7A**). Together, these findings indicate that OLs are relatively enriched in the outer membrane of both species under phosphate-limiting conditions, demonstrating that phosphate-responsive lipid remodeling is not uniformly distributed across the cell envelope and instead preferentially affects the membrane compartment most directly involved in environmental interactions.

**Figure 6.**
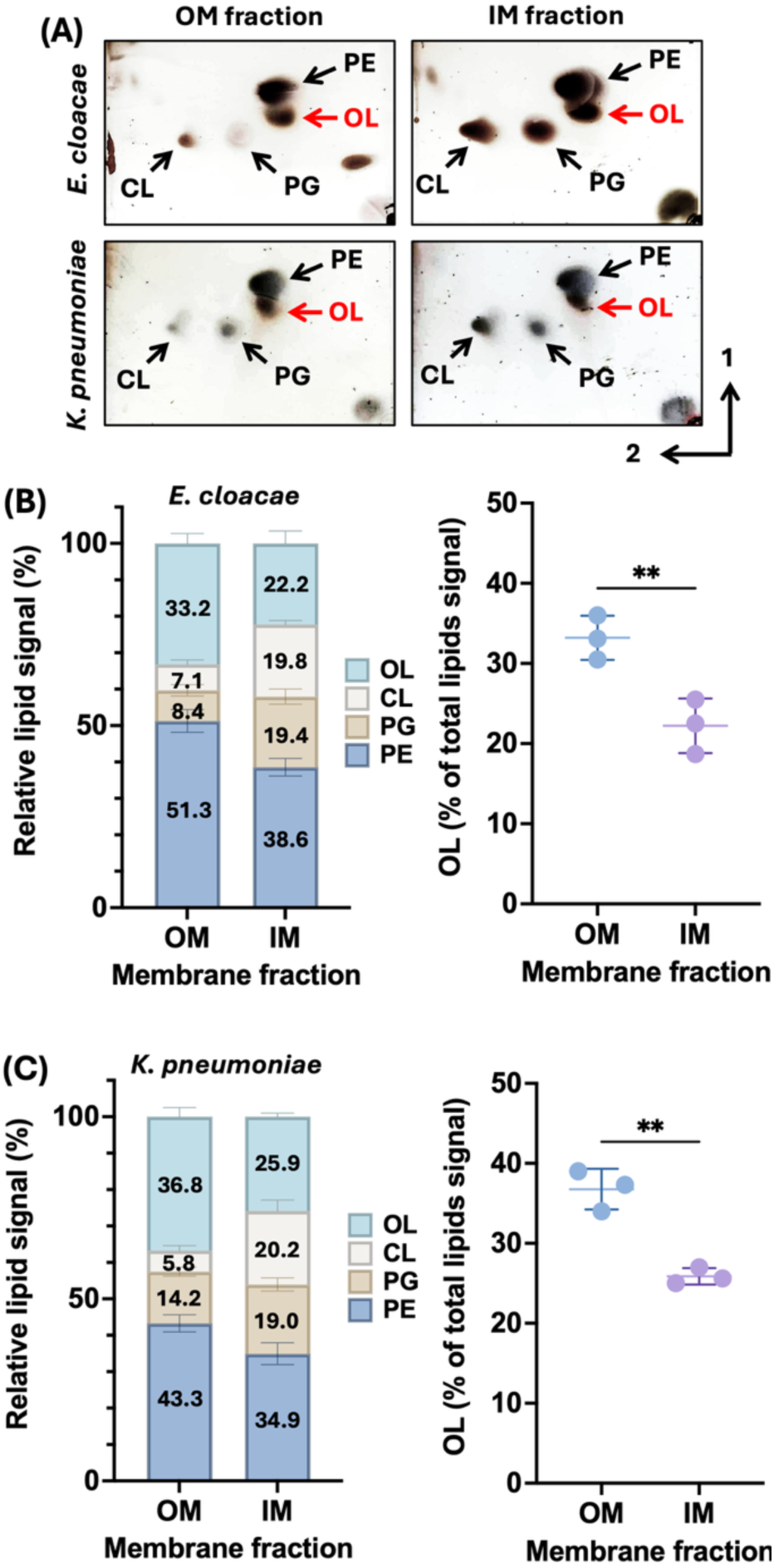
Ornithine lipids are enriched in the outer membrane of *E. cloacae* and *K. pneumoniae* during phosphate limitation. **(A)** Thin-layer chromatography (TLC) analysis of lipids extracted from outer membrane (OM) and inner membrane (IM) fractions of *E. cloacae* and *K. pneumoniae* grown under phosphate-limiting conditions (50 µM P_i_). Lipids were visualized by sulfuric acid staining. The positions of phosphatidylethanolamine (PE), phosphatidylglycerol (PG), cardiolipin (CL), and ornithine lipids (OL) are indicated. Representative TLC images are shown. **(B and C)** Relative lipid composition of OM and IM fractions from *E. cloacae* **(B)** and *K. pneumoniae* **(C)**, determined by densitometric analysis of TLC-separated lipids. Stacked bars represent the mean relative abundance of each lipid class, expressed as a percentage of the total lipid signal within each membrane fraction. OLs accounted for a greater proportion of the lipid signal in OM fractions than in IM fractions in both species. Individual data points represent independent biological replicates, and bars show the mean ± standard deviation (SD; *n* = 3). Statistical significance was assessed using a two-tailed paired Student’s *t* test. *P* < 0.01.

In *E. coli* W3110 expressing *E. cloacae olsF*, OLs were likewise detected in both inner and outer membrane fractions (**Figure S7B**), demonstrating that heterologously produced OLs are distributed across both membrane compartments. These findings suggest that outer membrane localization of OLs does not depend on species-specific features unique to *E. cloacae* or *K. pneumoniae*. Collectively, these results identify preferential outer membrane accumulation as a defining feature of OlsF-dependent lipid remodeling and provide a spatial framework for understanding how ornithine lipids influence envelope physiology and polymyxin tolerance.

## Discussion

The bacterial cell envelope must remain functional as nutrient availability fluctuates, even when nutrient limitation constrains membrane biogenesis. Here, we characterize a phosphate-responsive membrane adaptation in *E. cloacae* and *K. pneumoniae* in which phosphate limitation promotes OlsF-mediated OL synthesis. Our findings support a role for PhoBR in regulating OL accumulation in *E. cloacae* and show that OlsF-derived OLs are present in both membrane compartments, with OLs representing a greater proportion of the lipid profile in the outer membrane. We further demonstrate that OL production contributes to polymyxin tolerance, thereby linking nutrient availability to membrane lipid remodeling and antimicrobial susceptibility. Collectively, these findings identify OlsF-dependent OL synthesis as a phosphate-responsive envelope adaptation pathway in clinically relevant Enterobacterales.

OL biosynthesis is widespread among Gram-negative bacteria, although its genetic organization and distribution vary across bacterial lineages. Our focused analysis of recently reported genomic datasets revealed that OlsF is broadly but non-uniformly distributed among Enterobacterales. OlsF homologs were detected throughout the examined *Enterobacter* and *Klebsiella* reference genomes and were prevalent in several other genera, whereas no OlsF, OlsB, or OlsH homologs meeting our criteria were identified among the examined *Escherichia*, *Shigella*, *Salmonella*, or *Yersinia* genomes. Thus, OlsF-dependent OL synthesis appears to be conserved in certain Enterobacterales lineages rather than being a universal feature of the order, suggesting that OL-mediated membrane remodeling may represent a shared adaptive strategy among clinically important members of this group. Unlike the canonical OlsB/OlsA pathway, OlsF combines both acyltransferase activities within a single polypeptide (16). The conservation of H83 and H349 among OlsF homologs from *Enterobacter*, *Klebsiella*, and *Serratia*, together with the loss of detectable OL production following substitution of either residue, supports their importance for OlsF function.

Despite these differences in biosynthetic organization, phosphate-responsive OL production appears to be linked to the PhoBR regulatory system in several Gram-negative bacteria. In *E. cloacae*, disruption of *phoR* abolished detectable OL accumulation, consistent with PhoBR-dependent regulation of this pathway. Previous studies have also implicated PhoBR in OL biosynthesis in *P. aeruginosa* and *A. baumannii*, where OL production involves the separate OlsB and OlsA enzymes (9, 15). In *A. baumannii*, for example, *olsB* responds to phosphate limitation even though *olsA* is located at a separate chromosomal locus. These observations indicate that phosphate-responsive regulation of OL biosynthesis occurs across different genetic architectures. Together, these observations suggest that phosphate-responsive OL production has emerged across distinct biosynthetic architectures, although its regulation remains dependent on the physiological context of each organism (10, 12, 20). The repeated association between phosphate sensing and OL production nevertheless suggests that phosphorus-responsive membrane remodeling represents a broadly conserved adaptive strategy among diverse Gram-negative bacteria.

An important consequence of OlsF-dependent OL production is its contribution to polymyxin tolerance. Phosphate limitation increased polymyxin tolerance in both *E. cloacae* and *K. pneumoniae*, whereas deletion of *olsF* reduced the survival of *E. cloacae* following polymyxin B exposure. Conversely, heterologous expression of *E. cloacae olsF* increased tolerance to polymyxin B and colistin in *E. coli*, while OlsF variants that failed to produce detectable OLs did not confer the same increase in tolerance. Together, these loss- and gain-of-function experiments support a role for OL production in this phenotype. Previous studies have also linked OL accumulation to reduced polymyxin susceptibility. Early work in *Pseudomonas fluorescens* associated phosphate-induced OL production with increased polymyxin B resistance and reduced antibiotic binding (25). In *P. aeruginosa*, increased OL production through *olsBA* overexpression was associated with altered cell surface properties and reduced antimicrobial susceptibility (7), although an earlier study found that OL biosynthesis was not required for phosphate limitation-induced polymyxin resistance (15). More recently, OlsB-dependent aminolipid production was shown to promote colistin tolerance in *A. baumannii* (9), while OL biosynthesis contributed to polymyxin B tolerance in *V. cholerae* under low-salinity conditions (18). In addition, phosphate-induced glycolipid synthesis promotes polymyxin B tolerance in *P. aeruginosa* (8). Together, these findings indicate that the contribution of phosphorus-free membrane lipids to polymyxin susceptibility depends on the lipid biosynthetic pathway and physiological context. More broadly, multiple studies have demonstrated that remodeling of the Gram-negative cell envelope can alter susceptibility to antimicrobial stress and contribute to antibiotic adaptation (23, 26, 27).

How OL production contributes to polymyxin tolerance remains unclear. The accumulation of phosphorus-free OLs may alter the physicochemical properties of the envelope and potentially reduce the cellular requirement for phosphorus-containing phospholipids (2, 7). In *E. cloacae* and *K. pneumoniae*, OL production under phosphate limitation was accompanied by changes in the relative composition of membrane phospholipids. One possible explanation is that OL accumulation changes the physicochemical properties of the bacterial envelope, potentially influencing its interactions with polymyxins while reducing the phosphorus requirement for lipid synthesis. Under phosphate limitation, the accumulation of phosphorus-free OLs, accompanied by changes in relative phospholipid composition, may reduce the phosphorus requirement for membrane biogenesis. Because OLs lack a phosphate-containing headgroup, their accumulation could alter the distribution of charged lipid species within the bacterial envelope, potentially influencing electrostatic interactions with polycationic polymyxins. Although direct measurements of membrane surface charge and polymyxin binding will be required to evaluate this model, our findings suggest that nutrient-dependent lipid remodeling may simultaneously conserve phosphorus and modify envelope properties relevant to antibiotic susceptibility. Phosphate limitation can also promote polymyxin resistance through other envelope adaptations, including PmrAB-dependent lipid A modification, as recently reported in *E. coli* and *Salmonella* (28). Thus, OlsF-dependent OL production may be one component of a broader envelope response to phosphate limitation. Likewise, alterations in envelope architecture have been shown to facilitate adaptation to colistin exposure in other Gram-negative bacteria (27).

An unexpected finding of this study was the preferential enrichment of OlsF-derived OLs within the outer membrane. In *E. cloacae* and *K. pneumoniae*, OLs represented a greater proportion of the lipid profile in the outer membrane than in the inner membrane. This spatial organization is notable because the outer membrane is the primary interface with environmental stressors and serves as the initial site of interaction with polymyxins. Previous studies have also reported OL localization in the outer membrane of other Gram-negative bacteria, including *Rhizobium tropici* and *Ruegeria pomeroyi* (29, 30). OLs were also detected in both membrane fractions of *E. coli* expressing *E. cloacae olsF*, indicating that their distribution is not restricted to the native hosts. Their presence in the outer membrane is notable given that this compartment is the initial site of polymyxin interaction. How OLs reach the outer membrane, how their distribution is regulated, and whether they occupy its inner or outer leaflet remain unknown.

Beyond laboratory conditions, nutrient-responsive membrane remodeling may be relevant in host-associated environments, where phosphate availability fluctuates and bacterial pathogens encounter multiple envelope stresses. Recent studies have demonstrated that the physiological contributions of envelope maintenance pathways can be strongly influenced by environmental context, highlighting the importance of examining membrane adaptation under diverse stress conditions (31).

In *Burkholderia cenocepacia*, the production of phosphorus-free glycolipids was recently shown to contribute to intracellular survival in macrophages and virulence in an insect infection model (32). These findings illustrate how phosphate-responsive lipid remodeling can have consequences beyond phosphorus conservation. Our results raise the possibility that OL production similarly supports adaptation to host-associated conditions, although its contribution during *E. cloacae* or *K. pneumoniae* infection remains unknown.

Together, our findings establish OlsF-dependent OL biosynthesis as part of the membrane response to phosphate limitation in *E. cloacae* and *K. pneumoniae*. We propose a model (**Figure S8**) in which phosphate limitation promotes OL production through a response involving PhoBR, with OlsF-derived OLs present in both membrane compartments and representing a greater proportion of the lipid profile in the outer membrane. By contributing to polymyxin tolerance, OlsF-mediated lipid remodeling links phosphate availability to outer membrane composition and antimicrobial susceptibility, highlighting how nutrient-responsive lipid adaptations influence the functional properties of the Gram-negative cell envelope.

## Acknowledgements

The work was supported by funding from the National Institutes of Health (grants R35GM143053 to J.M.B, R01AI178692 to K.L.P, and R35139659 to J.S.B.) and Welch Foundation F-1155-20250403 to J.S.B. The funders had no role in study design, data collection and analysis, decision to publish, or preparation of the manuscript. All mass spectrometry data related to identification of lipids is available in the following repository: ID: MassIVE MSV000103672.

## Materials and Methods

### Bacterial strains and growth conditions

All bacterial strains and plasmids used in this study are listed in **Table S5**. *E. cloacae*, *K. pneumoniae*, *P. aeruginosa*, and *E. coli* strains were routinely recovered from frozen stocks on Luria-Bertani (LB Miller) agar at 37 °C. Isolated colonies were used to inoculate LB Miller broth or Tris minimal succinate (TMS) medium (8, 9), as indicated for each experiment. When required for plasmid maintenance or strain construction, antibiotics were added at the following concentrations: kanamycin, 25 µg mL⁻¹; chloramphenicol, 40 µg mL⁻¹; ampicillin, 100 µg mL⁻¹; and hygromycin, 250 µg mL⁻¹ for *E. coli* and 200 µg mL⁻¹ for *E. cloacae*.

TMS medium contained 20 mM sodium succinate, 200 mg L⁻¹ NaCl, 450 mg L⁻¹ NH₄Cl, 200 mg L⁻¹ CaCl₂, 200 mg L⁻¹ KCl, 450 mg L⁻¹ MgCl₂, 10 mg L⁻¹ FeCl₂, 10 mg L⁻¹ MnCl₂, and 10 mM HEPES (pH 7). Na₂HPO₄ was added to achieve the indicated final phosphate concentration. Phosphate-replete and phosphate-limiting conditions contained 1 mM and 50 µM P_i_, respectively, whereas 400 µM P_i_ was used where indicated for *P. aeruginosa*. All medium components were prepared in deionized water and sterilized by filtration through 0.22-µm pore-size filters.

For phosphate-dependent experiments, cultures were maintained at the indicated phosphate concentration throughout growth, including overnight cultures used to initiate experimental cultures, thereby minimizing phosphate carryover between conditions. Recombinant *E. coli* W3110 strains carrying *olsF* expression plasmids or the corresponding empty vector were grown in LB Miller supplemented with the appropriate selective antibiotic, 1 mM IPTG for induction of gene expression, and 1 mM L-ornithine. Unless otherwise indicated, cultures were incubated at 37 °C with shaking at 225 rpm.

### Construction of *E. cloacae* deletion mutants

The primers used in this study are listed in **Table S6**. Gene deletion mutants of *E. cloacae* were generated using a previously described λ-Red recombineering approach, with modifications (33–35). Briefly, pACBSR-Hyg was used to provide the λ-Red recombination machinery, and an FRT-flanked kanamycin resistance cassette was amplified from pKD4 using primers containing 75 and 125-nt extensions corresponding to the chromosomal regions flanking the target locus.

*E. cloacae* carrying pACBSR-Hyg was grown at 30 °C in LB Miller without added NaCl and supplemented with 200 µg mL⁻¹ hygromycin. Expression of the λ-Red recombination system was induced with 100 mM L-arabinose, and cultures were grown to an OD_600_ of approximately 0.4. Cells were then washed six times with ice-cold 10% glycerol and electroporated with 5 µg of the purified PCR product in a 2-mm cuvette at 1.8 kV. Following electroporation, cells were recovered for 4 h in LB Miller without added NaCl supplemented with 100 mM L-arabinose and plated on LB agar containing 35 µg mL⁻¹ kanamycin. Mutants were identified and confirmed by PCR.

To generate unmarked deletion mutants, the FRT-flanked kanamycin resistance cassette was removed using pCP20 as previously described (36). Mutants carrying the kanamycin cassette were transformed with pCP20 and selected on LB agar containing 100 µg mL⁻¹ ampicillin and 40 µg mL⁻¹ chloramphenicol at 30 °C. Transformants were subsequently grown in LB without antibiotic selection at 43 °C to promote FLP-mediated cassette excision and loss of pCP20. Colonies were screened for sensitivity to kanamycin and ampicillin, and cassette excision was confirmed by PCR.

Finally, pACBSR-Hyg was cured by serial passage of the mutant strains in LB without antibiotic selection. Colonies were screened for loss of hygromycin resistance, and hygromycin-sensitive isolates lacking the kanamycin resistance cassette were retained for subsequent experiments.

### Plasmid construction and complementation

The primers used for plasmid construction are listed in **Table S6**. The *olsF* coding sequences from *E. cloacae* and *K. pneumoniae*, as well as *phoR* from *E. cloacae*, were amplified by PCR and cloned into pMMB67EH-KanR. The *E. cloacae olsF* coding sequence was cloned into pMMB67EH-KanR using the SacI and SalI restriction sites, whereas *K. pneumoniae olsF* and *E. cloacae phoR* were cloned using EcoRI and HindIII. The *E. cloacae olsF* coding sequence was also cloned into pJM-HygR using the NcoI and HindIII restriction sites.

To assess the importance of conserved histidine residues in OlsF, H83 and H349 of *E. cloacae* OlsF were individually replaced with alanine to generate the H83A and H349A variants. Corresponding mutant *olsF* alleles were synthesized by Twist Bioscience and cloned into both pMMB67EHKanR and pJM-HygR. The pMMB67EHKanR constructs were generated using SacI and SalI, whereas the pJM-HygR constructs were generated using NcoI and HindIII. All plasmid constructs were verified by sequencing.

The pMMB67EHKanR- and pJM-HygR-based *olsF* constructs were introduced into *E. coli* W3110 for heterologous expression experiments and into the *E. cloacae* Δ*olsF* mutant for genetic complementation. Comparable phenotypes were obtained with both plasmid systems in heterologous expression and complementation assays, indicating that the observed effects were not vector-specific. The pMMB67EHKanR-*phoR* construct was introduced into the *E. cloacae* Δ*phoR* mutant for complementation. Strains carrying pMMB67EHKanR-derived plasmids or the corresponding empty vector were grown with 25 µg mL⁻¹ kanamycin. Strains carrying pJM-HygR-derived plasmids or the corresponding empty vector were grown with hygromycin at 250 µg mL⁻¹ for *E. coli* and 200 µg mL⁻¹ for *E. cloacae*. Expression from both pMMB67EHKanR- and pJM-HygR-derived constructs was induced with 1 mM IPTG. For heterologous OlsF expression experiments in *E. coli* W3110, cultures were additionally supplemented with 1mM L-ornithine.

### Analysis of total lipids and aminolipids

Overnight cultures were diluted to an initial OD_600_ of approximately 0.05 in fresh TMS medium containing the corresponding phosphate concentration. Cultures were grown in 20 mL of TMS containing 1 mM P_i_ or 40 mL containing 50 µM or 400 µM P_i_ and incubated for 24 h at 37 °C to obtain comparable biomass for lipid analysis. Cells were collected by centrifugation, and total lipids were extracted using the Bligh and Dyer method, as previously described (37).

Equivalent amounts of total lipid extracts were applied to high-performance thin-layer chromatography (HPTLC) on silica gel plates and separated. For one-dimensional HPTLC analysis, plates were developed in chloroform:methanol:water (130:50:8, v/v/v). For two-dimensional HPTLC analysis, plates were developed in chloroform:methanol:water (140:60:10, v/v/v) in the first dimension and chloroform:methanol:acetic acid (130:50:20, v/v/v) in the second dimension.

Total lipids were visualized with 10% sulfuric acid in ethanol followed by heating at 150 °C. Aminolipids were detected with 0.2% ninhydrin in acetone followed by heating at 100 °C. Lipid spots were quantified by densitometry using ImageJ following background subtraction. The relative abundance of each lipid species was expressed as the percentage of its densitometric signal relative to the total lipid signal detected within the corresponding sample. These values represent relative staining intensities rather than absolute molar proportions. Quantitative values are reported in **Table S3**. Experiments were performed using three independent biological replicates. Representative HPTLC images are shown, and quantitative analyses were derived from all biological replicates.

### Mass spectrometry analysis

The dried lipid extracts from TLC bands obtained from Bligh and Dyer extractions of *E. cloacae* and *K. pneumoniae* were resuspended in 200 µL of chloroform. The resulting stock solution was subsequently diluted with a 1:1 (v/v) methanol: isopropanol for direct infusion. Shotgun lipidomic ESI-MS/MS analysis was conducted in negative-ionization mode using a Thermo Scientific Orbitrap Fusion Lumos Tribrid mass spectrometer. Nanoelectrospray ionization was performed using Pd/Au-coated, borosilicate static tips (OD 1.2 mm, ID 0.69 mm) pulled in-house at a spray voltage of −1000 V and a capillary temperature of 275 °C. Mass spectra were acquired at a resolving power of 50,000 (at *m/z* 200), with an automatic gain control (AGC) target of 5E4 and an RF lens setting of 60%. All MS1 and MS/MS spectra were averaged over 50 scans. The ten most abundant precursor ions were selected for higher-energy collisional dissociation (HCD) for lipid identification. MS/MS spectra were acquired using a precursor isolation width of 0.7 *m/z* and normalized collision energies (NCEs) ranging from 15% to 27%. Data was manually interpreted using Thermo Xcaliber Qual Browser and ChemDraw. Precursor and fragment ions were assigned within 10 ppm mass error.

### Microscopy and image analysis

For morphological analysis, mid-exponential-phase cultures (OD₆₀₀ = 0.4–0.5) were labeled with 100 µM HADA (7-hydroxycoumarin-3-carboxylic acid–D-alanine) for 15 min to visualize active peptidoglycan synthesis (31). Cells were fixed with 1.6% paraformaldehyde, washed with PBS, and mounted on 1.5% agarose pads prepared in PBS. Images were acquired using a Nikon Eclipse Ti-2 wide-field epifluorescence microscope equipped with a Plan Apo 100×/1.45 NA objective and a Photometrics Prime 95B camera using NIS-Elements software. Cell length was quantified using ImageJ/Fiji with the MicrobeJ plugin (38). At least *n* = 150 cells were analyzed per condition from three independent biological replicates

### Polymyxin susceptibility and survival assays

Polymyxin susceptibility was assessed using inhibition zone and survival assays. For inhibition zone assays, Petri dishes containing 20 mL of agar prepared with the corresponding phosphate concentration were overlaid with 5 mL of soft agar containing approximately 10⁸ cells. After solidification, sterile 6-mm filter-paper disks were placed on the agar surface, and 10 µL of 2 mg mL⁻¹ polymyxin B or colistin was applied to each disk. Plates were incubated at 37 °C for 24 h, after which inhibition zone diameters were measured. For survival assays, overnight cultures were grown in 5 mL of the corresponding medium in 18 × 150 mm glass culture tubes, and all polymyxin exposures were performed at 37 °C in the same culture volume. *K. pneumoniae* and *E. coli* W3110 cultures were incubated with shaking at 225 rpm, whereas *E. cloacae* cultures were incubated statically. For phosphate-dependent survival assays, *E. cloacae* and *K. pneumoniae* cultures were adjusted to an OD₆₀₀ of 0.150 in 5 mL of fresh medium containing either 1 mM or 50 µM Pi and exposed to 8 µg mL⁻¹ polymyxin B. Samples were collected at 0, 3, and 6 h, serially diluted, and spotted onto LB agar to assess survival following antibiotic exposure.

For complementation assays, overnight cultures of *E. cloacae* strains carrying the o*lsF* expression plasmid or the corresponding empty vector were grown in the presence of the appropriate selective antibiotic and 1 mM IPTG. The following day, cells were collected by centrifugation and washed once with fresh phosphate-limiting medium. Washed cells were adjusted to an OD₆₀₀ of 0.150 in 5 mL of fresh phosphate-limiting medium containing 1 mM IPTG but no selective antibiotic and exposed to 24 µg mL⁻¹ polymyxin B. Samples were collected at 0, 3, and 6 h, serially diluted, and spotted onto LB agar.

For heterologous expression assays, overnight cultures of *E. coli* W3110 carrying the *E. cloacae olsF* expression plasmid, the corresponding OlsF variants, or the empty vector were grown in LB containing the appropriate selective antibiotic, 1 mM IPTG, and 1 mM L-ornithine. The following day, cells were collected, washed once with fresh LB, and resuspended in 5 mL of fresh LB to a final OD₆₀₀ of 0.2. The medium was supplemented with 1 mM IPTG and 1 mM L-ornithine, without selective antibiotics. Polymyxin B or colistin was added to a final concentration of 1 µg mL⁻¹. Cultures were incubated at 37 °C with shaking at 225 rpm, and samples were collected at 0, 30, and 60 min, serially diluted, and spotted onto LB agar.

Survival assays were performed under the specified experimental conditions, with polymyxin concentrations differing among assays involving phosphate availability, *olsF* complementation, and heterologous expression in *E. coli* W3110.

### Inner and outer membrane fractionation

Inner and outer membranes were separated by discontinuous sucrose density-gradient ultracentrifugation as previously described (39), with modifications. Overnight cultures were diluted to an initial OD_600_ of 0.01 in fresh medium and grown for 16 h at 37 °C. *E. cloacae* and *K. pneumoniae* were grown in 1 L of TMS medium containing 50 µM P_i_, whereas *E. coli* W3110 carrying the *E. cloacae olsF* expression plasmid was grown in 400 mL of LB supplemented with 25 µg mL⁻¹ kanamycin, 1 mM IPTG, and 1 mM L-ornithine. For *K. pneumoniae*, 0.5 µM EDTA was added to the culture medium to facilitate cell pelleting, as previously described (34). Culture volumes were adjusted empirically to obtain sufficient membrane material for fractionation and downstream lipid analyses.

Cells were harvested by centrifugation and resuspended in 12.5 mL of 0.5 M sucrose and 10 mM Tris-HCl (pH 7.5). Lysozyme was added to a final concentration of 144 µg mL⁻¹, and samples were stirred on ice for 2 min. An equal volume (12.5 mL) of 1.5 mM EDTA was then added, and samples were stirred for an additional 7 min on ice.

Cells were collected by centrifugation and resuspended in 25 mL of 0.2 M sucrose and 10 mM Tris-HCl (pH 7.5) containing 55 µL of 1 M MgCl₂, 1 µL of RNase/DNase, and 1 µL of protease inhibitor cocktail. Cells were disrupted by six passages through an EmulsiFlex high-pressure homogenizer. Unbroken cells and cellular debris were removed by centrifugation, and total membranes were collected by ultracentrifugation at 184,500 × g for at least 1 hour at 4 °C.

Total membrane pellets were resuspended in 1 mL of 20% (w/v) sucrose containing 1 mM EDTA and 1 mM Tris-HCl (pH 7.5). Discontinuous sucrose gradients were prepared with 2 mL of 73% (w/v) sucrose and 5 mL of 53% (w/v) sucrose, each containing 1 mM EDTA and 1 mM Tris-HCl (pH 7.5). The total membrane fraction was layered above the 53% sucrose solution, and the remaining volume was filled with 20% sucrose solution. Gradients were centrifuged at 288,000 × g for 16 hours at 4 °C.

Membrane bands at the 20–53% and 53–73% sucrose interfaces were collected as the inner membrane and outer membrane fractions, respectively. Approximately 1 mL of each fraction was diluted with 9 mL of 10 mM Tris-HCl (pH 7.5) to remove sucrose. Membranes were collected by ultracentrifugation at 184,500 × g for 1 hour at 4 °C and resuspended in 200 µL of 10 mM Tris-HCl (pH 7.5). Protein concentrations were determined using the Bradford assay (40). The purity of membrane fractions was evaluated by measuring NADH oxidase activity and lipopolysaccharide enrichment as markers of the inner and outer membranes, respectively.

### Validation of membrane fractionation and lipid analysis

Membrane separation was assessed by NADH oxidation and LPS enrichment based on previously described methods (39). NADH oxidation assays were performed using 50 µg of membrane protein. Samples were brought to 990 µL with 10 mM Tris-HCl (pH 7.5), followed by addition of 10 µL of a 10 mg mL⁻¹ NADH solution. NADH oxidation was monitored by measuring the decrease in absorbance at 340 nm every 30 sec for 5 min. Enrichment of NADH-oxidizing activity was used as a marker of the inner membrane purity and enrichment.

For LPS analysis, inner membrane and outer membrane fractions were normalized to 50 µg of membrane protein, treated with Proteinase K, separated by SDS-PAGE, and stained using the Pro-Q Emerald 300 Lipopolysaccharide Gel Stain Kit (Thermo Fisher Scientific, P20495). LPS enrichment was used as a marker of outer membrane purity and enrichment.

Following removal of aliquots for NADH oxidation and LPS analyses, the remaining material from each IM and OM fraction was subjected to Bligh-Dyer lipid extraction. Lipid extracts were separated by HPTLC and visualized as described above. Lipid spots were quantified by densitometry using ImageJ, and OL abundance was expressed as a percentage of the total lipid signal detected within each membrane fraction. Membrane validation and lipid analyses were performed using three independent biological replicates.

### Comparative analysis of OlsF distribution and sequence conservation

The distribution of OlsF within Enterobacterales was assessed by reanalyzing the filtered BLASTp dataset and associated RefSeq taxonomy reported by Moyano-Palazuelo et al. (20), rather than performing an independent BLAST search. The original analysis used complete annotated bacterial reference genomes available in RefSeq and identified OlsF homologs by BLASTp using OlsF proteins from *S. proteamaculans* and *Flavobacterium johnsoniae* as queries. Hits were retained using the criteria reported in the original study: an E-value ≤ 1 × 10⁻¹⁰, ≥30% amino acid sequence identity, ≥70% query coverage, and ≥70% subject coverage. For the present analysis, reference genomes assigned to Enterobacterales were grouped by genus according to the associated RefSeq taxonomy, and the proportion of genomes containing an OlsF homolog was determined for each genus (**Figure 1A**; **Table S1**).

For sequence conservation analysis, OlsF protein sequences from *Enterobacter*, *Klebsiella*, and *Serratia* were obtained from the same dataset. Duplicate RefSeq protein accessions were removed, resulting in a dataset of 48 unique OlsF sequences. Protein sequences were aligned using Clustal Omega (41). Residue positions were referenced to OlsF from *S. proteamaculans* (WP_099062946.1), and conservation of the positions corresponding to H83 and H349 was determined from the multiple sequence alignment. Sequence logos spanning residues 76–90 and 342–356 of *S. proteamaculans* OlsF were generated from the aligned sequences using WebLogo 3 (42) (**Figure 1B**). These analyses were used to identify conserved residues for mutagenesis and to assess the prevalence of OlsF among representative Enterobacterales genomes.

